# In-depth proteomic analysis of *Plasmodium berghei* sporozoites using trapped ion mobility spectrometry with parallel accumulation-serial fragmentation

**DOI:** 10.1101/2020.11.26.400192

**Authors:** Soumia Hamada, Cédric Pionneau, Christophe Parizot, Olivier Silvie, Solenne Chardonnet, Carine Marinach

## Abstract

Malaria is caused by *Plasmodium* spp. protozoan parasites, which are transmitted by female anopheline mosquitoes in the form of sporozoites. Once deposited in the dermis during the blood meal of the mosquito, sporozoites rapidly migrate to the liver for an initial and obligatory round of replication inside hepatocytes, before exponential multiplication of the parasite in the blood and onset of the malaria disease. Sporozoites and liver stages provide attractive targets for the development of a malaria vaccine. Until now, a single antigen from *Plasmodium falciparum*, the deadliest species infecting humans, has been considered for subunit vaccine clinical development, with limited success so far. This emphazises the need to identify novel targets. In this context, defining the parasite proteome is important not only to guide the down-selection of potential candidate antigens, but also to allow a better understanding of the parasite biology. Previous studies have determined the total proteome of sporozoite stages from the two main human malaria parasites, *P. falciparum* and *P. vivax*, as well as *P. yoelii*, a parasite that infects rodents. Another murine malaria parasite, *P. berghei*, has been widely used to investigate the biology of *Plasmodium* pre-erythrocytic stages. However, a deep view of the proteome of *P. berghei* sporozoites is still missing. To fill this gap, we took advantage of a novel highly sensitive timsTOF PRO mass spectrometer, based on trapped ion mobility spectrometry with parallel accumulation-serial fragmentation. Combined with three alternative methods for sporozoite purification, this approach allowed us to identify the deep proteome of *P. berghei* sporozoites using low numbers of parasites. This study provides a reference proteome for *P. berghei* sporozoites, identifying a core set of proteins expressed accross species, and illustrates how the unprecedented sensitivity of the timsTOF PRO system enables deep proteomic analysis from limited sample amounts.

## 1. Introduction

Malaria, with more than 200 millions estimated cases and 400,000 deaths every year [1], remains a major public health problem in many countries. Significant progress has been achieved over the past decades in reducing malaria incidence and mortality, through the systematic use of insecticide-treated bednets and potent antimalarial artemisinin-based drug combinations. However, progress in malaria control has recently stalled, and the continued emergence of parasite and mosquito resistance to antimalarial medicines and insecticides, respectively, is a serious threat to malaria control. The disease is caused by parasites of the genus *Plasmodium*, which are transmitted via the bite of infected *Anopheles* female mosquitoes. Invasive forms called sporozoites present in the salivary glands of the vector are deposited in the dermis during a blood meal. After traversing the skin, motile sporozoites traffic to the liver through the blood stream, and reach the liver parenchyma where they invade hepatocytes for an initial and obligatory replication phase, resulting in the release of tens of thousands of merozoites [2]. These merozoites invade erythrocytes and initiate the exponential asexual reproduction of the parasite in erythrocytes, causing the symptomatic phase of malaria. Of the *Plasmodium* species infecting humans, *Plasmodium falciparum* is the most prevalent and the deadliest species, especially in subsaharan Africa. *P. vivax*, the second most important species in humans, is widely distributed around the world but causes less severe malaria. One particularity of *P. vivax* is to cause relapsing malaria episodes, due to hypnozoites, which are dormant liver-stage parasites that can reactivate weeks or months after parasite transmission by a mosquito.

Infection of the liver by sporozoites is an essential and clinically silent phase of the malaria life cycle, and has long been considered as an ideal target for a malaria vaccine [3]. However, despite intense research efforts, no efficacious malaria vaccine has been licensed yet. Until now, a single antigen, the circumsporozoite protein (CSP), has been pursued as a vaccine target against the extracellular sporozoite stage. The principal malaria vaccine candidate, RTS,S, targets CSP from *P. falciparum*, yet confers only partial and short-lived protection [4]. This emphasizes the need to develop more efficacious malaria vaccines targeting other antigens. In this context, proteomic studies of sporozoites, by identifying the set of proteins expressed by these forms, can guide the down-selection of potential new candidates.

Working with limited sample amounts has long been a challenge for in-depth proteome analysis. Studying the *Plasmodium* sporozoite proteome magnifies such a problem. Indeed, sporozoites can only be obtained from infected mosquitoes and must be isolated by hand dissection of the insect salivary glands. In addition, sporozoites are small cells (around 1 μm x 10-15 μm) that can be obtained only in limited numbers (usually less than 10^5^ per mosquito). Recovery of high numbers of sporozoites for downstream proteomic studies typically requires dissecting hundreds of mosquitoes, resulting in a massive contamination of the samples with mosquito material. Methods have been developed to purify sporozoites and limit the proportion of mosquito proteins [5]. These approaches, combined with LC-MS/MS, led to the characterization of the total proteome of *P. falciparum* and *P. vivax* salivary gland sporozoites, where the most comprehensive studies identified 2,039 and 1,972 proteins respectively (out of more than 5,500 proteins encoded by their genome) [6–10]. Similar studies performed in the rodent *P. yoelii* identified 1,774 proteins in salivary gland sporozoites of this species [8,10]. Surface proteomes were also described for the three species, based on chemical labeling of live parasites followed by LC-MS/MS [8,9,11].

*P. berghei* is another rodent malaria parasite that has been widely used as a model to study *Plasmodium* pre-erythrocytic stages. *P. berghei* is closely related to *P. yoelii* yet shows differences in the invasion routes used by sporozoites to infect hepatocytes. While *P. yoelii*, like *P. falciparum*, relies on the host protein CD81 to infect hepatocytes [12], *P. berghei* can use the scavenger receptor B1 (SR-B1) as an alternative entry route [13,14]. As SR-B1 is also involved during *P. vivax* infection [13], *P. berghei* provides a suitable model to study SR-B1-dependent liver infection [15]. The molecular basis of this differential usage of host receptors is not fully elucidated, but the sporozoite 6-cysteine domain protein P36 was shown to play a central role [13]. Although *P. berghei* is the most studied rodent parasite, the sporozoite proteome for this species has been solved only partially, with 134 proteins identified in one report [16]. Importantly, in the current version of PlasmoDB (v49) [17], proteins with reported mass spectrometry evidence in *P. berghei* sporozoites correspond to an extrapolation from proteomic studies performed with *P. yoelii* sporozoites [8].

Here, we took advantage of the highly sensitive timsTOF PRO mass spectrometer to determine the total proteome of *P. berghei* sporozoites. This system is equipped with the trapped ion mobility spectrometry (TIMS) technology for parallel accumulation serial fragmentation (PASEF). The dual TIMS technology enables ions trapping in the front section and ions separation according to their ion mobility in the rear section. Combined with rapid quadrupole switching and TOF detector, the timsTOF PRO enables the fragmentation of multiple simultaneously eluting precursor ions with a near 100% duty cycle. The timsTOF PRO offers speed and sensitivity gains of up to 10-fold compared to other mass spectrometry approaches [18]. Combined with three alternative methods for sporozoite purification, this system allowed us to identify the deep proteome of *P. berghei* sporozoites using unprecedented low numbers of parasites.

## 2. Results and discussion

### 2.1. Efficiency of sporozoite purification

*Plasmodium* sporozoites are obtained by hand dissection of the salivary glands of infected mosquitoes, resulting in sample contamination with proteins from the mosquito or its microbiota. Purification of sporozoites after dissection is a crucial step to reduce the quantity of contaminating proteins of mosquito origin. We compared three different methods for parasite purification: 1) the density gradient purification procedure, developped by Kennedy et al [5], used as a reference method 2) immunocapture of sporozoites using magnetic beads coupled to anti-CSP antibodies and 3) sorting of fluorescent sporozoites by flow cytometry. **Figure 1** illustrates the workflow that was used for the proteomic analysis of *P. berghei* sporozoites. In total, 4 independent sporozoite preparations were purified with the density gradient method, 2 using magnetic beads and 4 by flow cytometry, as summarized in **Table 1**. For each purification method, we assessed both the sporozoite recovery rate, i.e. the proportion of sporozoites recovered after purification, and the efficiency of the purification at the protein level. For this purpose, lysates of purified sporozoites were analyzed by LC-MS/MS to determine the proportion of parasite-versus mosquito-derived proteins.

**Figure 1.**
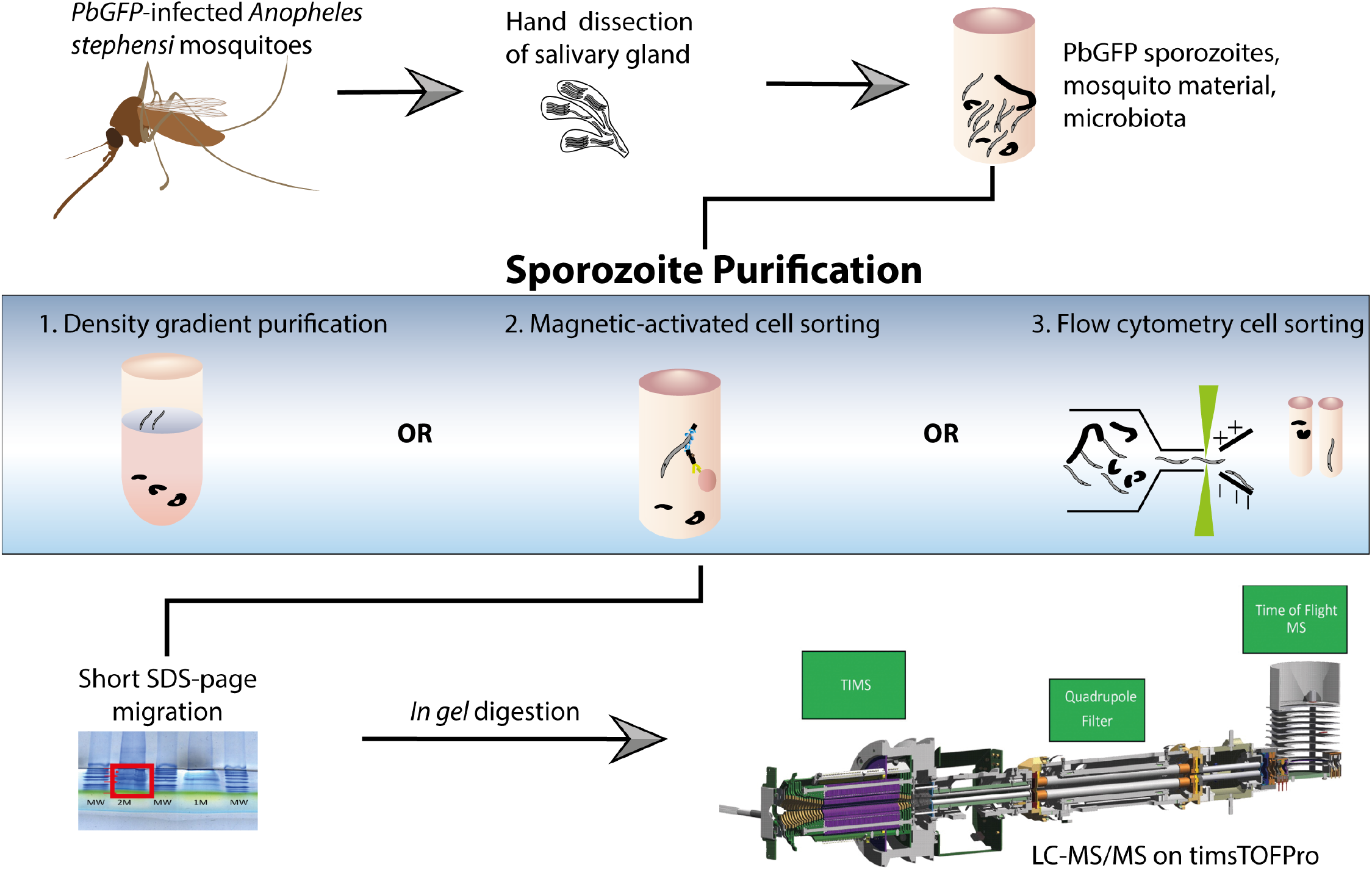
Workflow for sample preparation and proteomic analysis. GFP-expressing *P. berghei* sporozoites were collected by hand dissection of the salivary glands of infected *A. stephensi* mosquitoes, and purified either by density gradient centrifugation, immunocapture with magnetic beads or by flow cytometry cell sorting. Lysates from purified sporozoites were processed by SDS-PAGE and in-gel trypsin digestion, and resulting peptide samples were analyzed by LC-MS/MS with the TimsTOF Pro system.

**Table 1.**
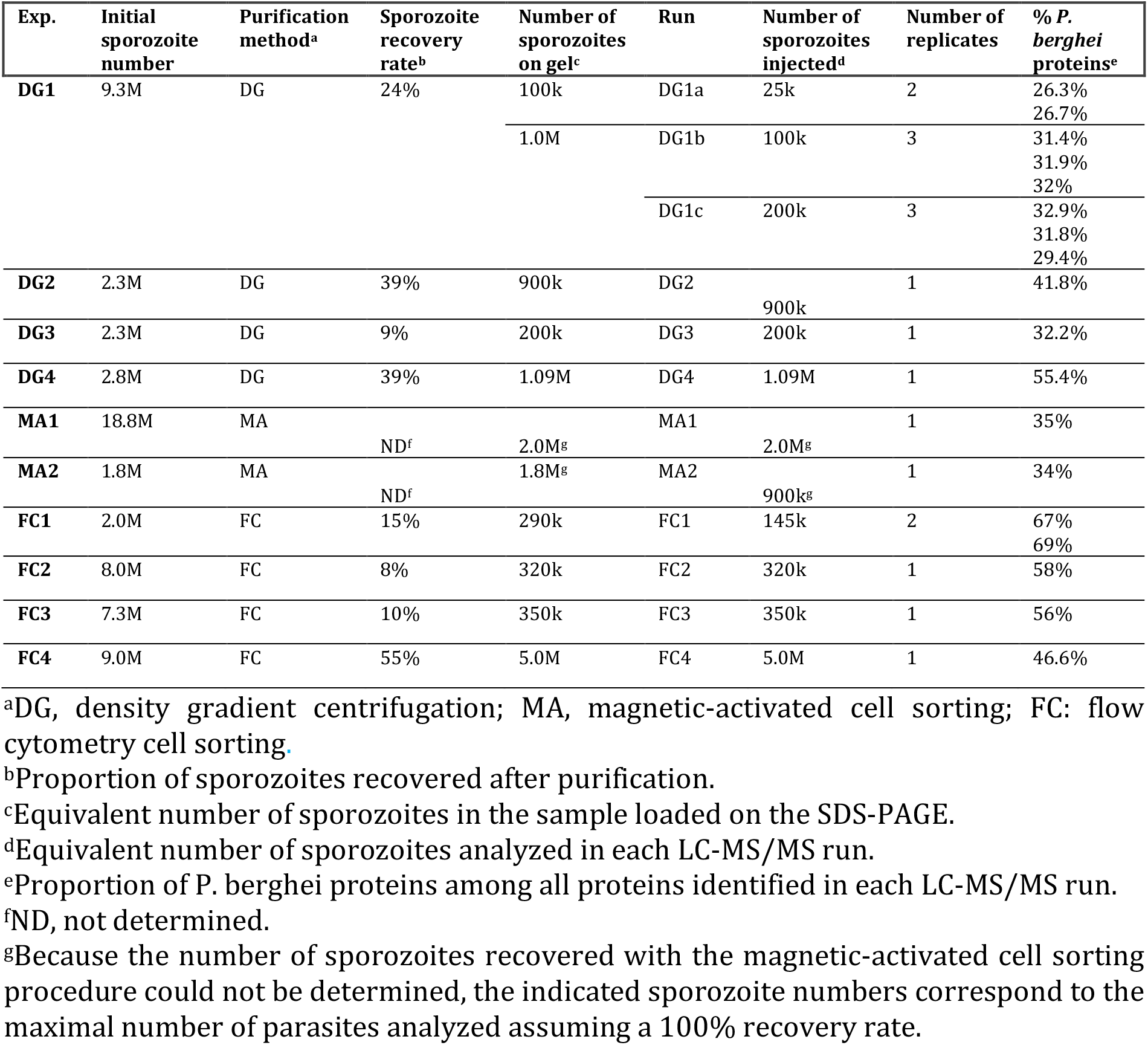
Summary of the samples analyzed in this study.

Purification using the density gradient protocol was relatively easy to execute but the sporozoite recovery rate was highly variable, ranging from 9 to 39% (mean 28%) (**Table 1**). Despite this lack of reproducibility in our hands, the purification efficiency was satisfying, with *P. berghei* proteins representing between 30 and 55% (mean 39.93%) of the total number of proteins identified by mass spectrometry (**Figure 2A**), which is consistent with values reported with other species [8]. Immunocapture of sporozoites with anti-CSP antibodies coupled to magnetic beads was easy and rapid to execute. However, mosquito debris remained abundant despite extensive bead washes. In addition, as sporozoites were agglutinated with the beads after elution, it was impossible to the determine the yield of recovered sporozoites with this method. Nevertheless, samples were analyzed by mass spectrometry, resulting in around 35% of identified proteins being of parasite origin (**Table 1** and **Figure 2A**). Sorting of GFP-expressing sporozoites by flow cytometry also showed highly variable recovery rates, varying between 10 and 55% (**Table 1**). The poor recovery rate observed with some of the samples is probably due to the intrinsic low efficiency of the sorting method and to the fact that sorted parasites were highly diluted during the procedure, increasing the risk of parasite loss during subsequent centrifugations. Nevertheless, the purification efficiency was satisfying, with *P. berghei* proteins representing up to 60% of the total number of proteins identified by mass spectrometry (**Table 1** and **Figure 2A**). Based on these results, we conclude that both immunocapture and FACS sorting of sporozoites provide valuable alternative to the density gradient reference method.

**Figure 2.**
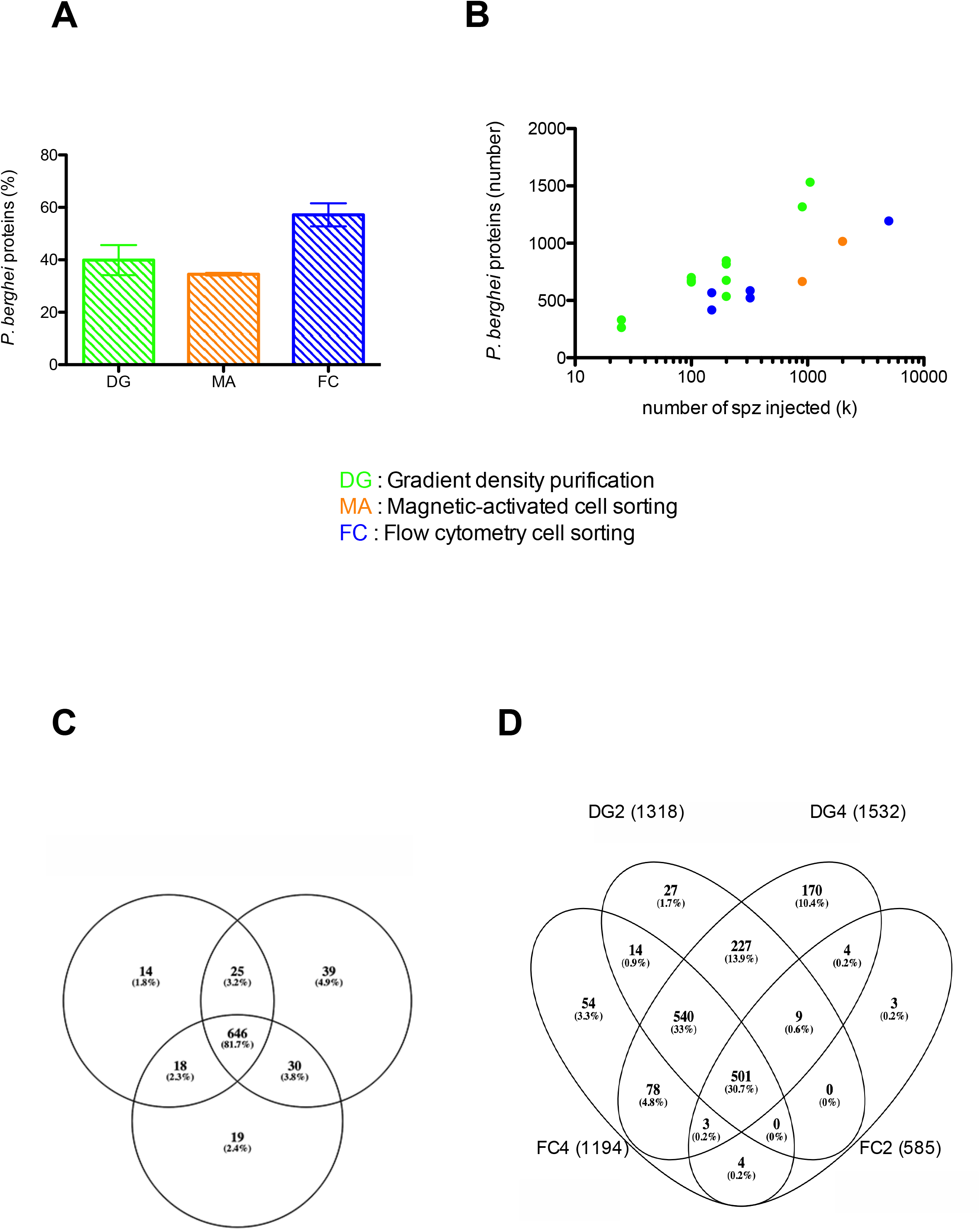
Effect of parasite purification and sample size on proteome identification. A. The efficiency of sporozoite purification was evaluated at the protein level by determining the proportion of *P. berghei* proteins among all proteins identified by mass spectrometry. The data are displayed as the mean (+/- SEM) from 2-4 experiments for each purification method. B. Number of *P. berghei* proteins identified by mass spectrometry depending on the sample size, represented as the equivalent number of sporozoites analyzed by mass spectrometry. Each data point corresponds to one MS/MS run. DG, density gradient centrifugation (green); MA, magnetic activated cell sorting (orange); FC, flow cytometry cell sorting (blue). C. Venn diagram showing the number of *P. berghei* proteins identified in 3 technical replicates of the DG1a sample analysis (100,000 sporozoites). D. Venn diagram showing the number of *P. berghei* proteins identified in 4 biological samples, each analyzed in a single run.

### 2.2. Effect of sporozoite sample size on proteome identification

We next analyzed the effect of sample size on the performance of the timsTOF PRO system for protein identification, for the three purification methods. Remarkably, mass spectrometry analysis of as few as 100,000-200,000 sporozoites purified by the density gradient method (DG1 and DG3 samples in **Table 1**) identified 500-900 *P. berghei* proteins (Figure 2B), and a single analysis of 900,000 sporozoites (DG2 sample in Table 1) resulted in the identification of 1532 *P. berghei* proteins (Figure 2B), which represents 93 % of the total number of proteins identified in the entire study (see below).

When combining all the identification results from 10 independent experiments (18 injections), we detected a total of 1648 proteins in *P. berghei* salivary gland sporozoites (Table S1 and Table S2). Increasing the number of technical replicates (multiple injections of the same sample) had a modest impact on the number of proteins identified (Figure 2C), with an overlap above 80%, which is more than previously reported on Orbitrap systems (59 to 76%) [19,20], and can probably be explained by the high frequency of the selection/fragmentation cycle (up to 100 per cycle). In contrast, increasing the number of biological independent samples increased substantially the number of proteins identified in *P. berghei* sporozoites (Figure 2D), with an overlapp of only 30,7%. The combined analysis of 10 samples (18 injections in **Table 1**) led to the identification of the 1648 *P. berghei* proteins listed in **Table S2**. The number of proteins identified in *P. berghei* sporozoites is within the same range as those reported in previous proteomic studies of *P. falciparum, P. vivax* and *P. yoelii* sporozoites (Figure 3A). However, 0.4-1×10^7^ purified salivary gland sporozoites were used for MS experiments in these studies [8–10], i.e. ~10 times more than reported here. Altogether, these results illustrate that the high sensitivity of the timsTOF PRO mass spectrometer allows in-depth identification of the *Plasmodium* sporozoite proteome, using low numbers of parasites.

**Figure 3.**
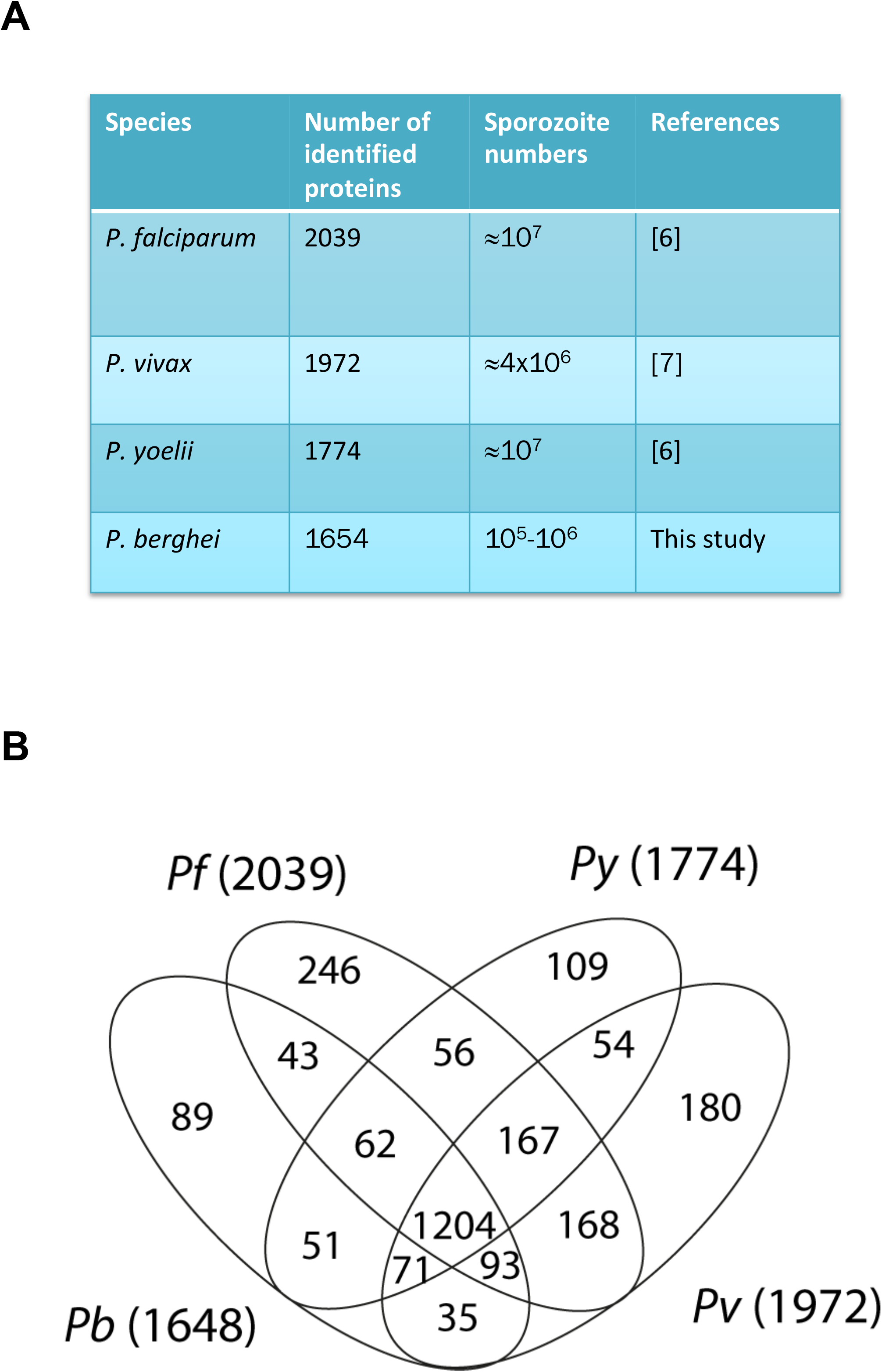
Comparison of the sporozoite proteomes across *Plasmodium* species. A. Summary of proteomic studies of *P. falciparum*, *P. vivax*, *P. yoelii* and *P. berghei* sporozoites, indicating the number of parasite proteins identified and the sample size. B. Venn diagram representation of the sporozoite proteome composition across *Plasmodium* species.

### 2.3. Identification of a core set of sporozoite proteins across *Plasmodium* species

Among the 1648 proteins identified in *P. berghei* sporozoites (**Table S2**), 1559 proteins (94.2%) have been identified in at least one other species (**Figure 3B** and **Table S3**). More precisely, 1402 proteins (84.8%) have been detected in *P. falciparum*, 1403 (84.9%) in *P. vivax* and 1388 (83.9%) in *P.yoelii*, showing overall extensive overlap. Comparison of the proteome datasets across species revealed a core set of 1204 proteins detected in salivary gland sporozoites from the four *Plasmodium* species (**Table S3**). In contrast, 51 (3.0%) proteins were identified only in sporozoites from the rodent malaria species (*P. yoelii* and *P. berghei*) and 89 (5.4%) only in *P. berghei*.

### 2.4. Analysis of sporozoite protein families

We next scrutinized the *P. berghei* proteome dataset for a selection of protein families, in comparison with published proteomes from *P. falciparum, P. vivax* and *P. yoelii* (**Table S3**). Like other apicomplexan invasive stages, sporozoites possess specialized apical secretory organelles termed micronemes and rhoptries, whose regulated secretion plays a key role during parasite locomotion, migration through the host tissues and invasion of hepatocytes. Overall, micronemal and rhoptry proteins were well conserved between sporozoite species. Many well-characterized micronemal proteins were shared across the sporozoite proteome datasets, including TRAP family members (TRAP, TREP), proteins involved in cell traversal (SPECT, PLP1, CelTOS and GEST), AMA1 and MAEBL, and members of the 6-cysteine domain (6-cys) protein family (P36, P52, P38 and B9). The claudin-like apicomplexan microneme protein (CLAMP), recently identified as a conserved apicomplexan protein that is essential for host cell invasion by *Toxoplasma gondii* tachyzoites and *P. falciparum* merozoites [21], was also identified in sporozoites from the four species, yet its role in this stage is still unknown. Eight rhoptry neck (RON) proteins were identified across sporozoite species (RON1/ASP, RON2, RON3, RON4, RON5, RON6, RON11 and RON12), as well as the rhoptry bulb high molecular weight rhoptry protein (RhopH) 2 and 3.

We also noted some differences between sporozoite proteomes. For example, the merozoite TRAP-like protein (MTRAP) was identified in *P. berghei*, *P. vivax* and *P. yoelii* but not in *P. falciparum* sporozoite samples. MTRAP is essential for gamete egress and formation of oocysts in the mosquito [22], but its role in sporozoites is unknown. The 6-cys protein P12p was detected in *P. berghei, P. yoelii* and *P. falciparum*, but not in *P. vivax*. Among rhoptry proteins, the Cytosolically Exposed Rhoptry Leaflet Interacting protein 1 (CERLI1), was identified in *P. berghei* and *P. vivax* sporozoites only. CERLI1 (also referred to as RASP2) plays an essential role in *P. falciparum* merozoites and *T. gondii* tachyzoites for rhoptry secretion and host cell invasion [23,24]. Subtilisin-like (SUB) proteins are essential serine proteases that cleave merozoite membrane proteins during invasion and egress [25,26]. Interestingly, SUB2 was identified in sporozoites from *P. falciparum, P. vivax* and *P. yoelii*, but not *P. berghei*. Conversely, SUB1 was detected in sporozoites from *P. berghei* but not in the other species. When looking at factors involved in gene regulation, we observed variations in the repertoire of AP2 transcription factors detected in the different sporozoite populations. While seven AP2 factors were detected in at least 3 species (including the previously characterized AP2-SP/EXP, AP2-I, AP2-L, AP2-O4 and AP2-O5), 4 members of the family were exclusively detected in *P. falciparum* sporozoites (including AP2-O), and one in *P. vivax* (AP2-O2). The sporozoite and liver stage asparagine-rich protein (SLARP), a master regulator of liver stage development [27,28], was identified in *P. falciparum*, *P. berghei* and *P. yoelii*, but not in *P. vivax*. Whether these differences reflect biological specificities or merely variations in protein expression and/or detection remains to be determined. Finally, 137 proteins among the 1205 core set (11%) correspond to proteins of unknown function.

## 3. Concluding remarks

This study provides a reference proteome dataset for *P. berghei* sporozoites, complementing previous proteomic studies of *P. falciparum*, *P. vivax* and *P. yoelii* parasites. We identify a conserved set of more than 1200 sporozoite proteins shared between the four species, and found some differences between the datasets. Further investigations will be required to determine whether these differences reflect biological specificities or merely variations in protein abundance. Finally, identification of the *P. berghei* sporozoite proteome was achieved with much lower numbers of cells as compared to previous studies. This illustrates how the high sensitivity of the timsTOF PRO system enables deep proteomic analysis from limited sample amounts. In the context of malaria, this opens novel perspectives to explore the parasite proteome, including in elusive stages such as the hypnozoites, or for proteogenomic studies of field isolates.

## 4. Experimental Section

### Ethics statement

All animal work was conducted in strict accordance with the Directive 2010/63/EU of the European Parliament and Council ‘On the protection of animals used for scientific purposes’. Protocols were approved by the Ethical Committee Charles Darwin N°005 (project #7475).

### Production of Plasmodium sporozoite-infected mosquitoes

We used GFP-expressing *P. berghei* (PbGFP, ANKA strain) parasites, obtained after integration of a GFP expression cassette at the dispensable p230p locus [29]. PbGFP blood stage parasites were propagated in female Swiss mice (6-8 weeks old, from Janvier Labs). *Anopheles stephensi* mosquitoes were fed on PbGFP-infected mice using standard methods, and kept at 21°C. Infected mosquitoes were fed on an 10% w/v sucrose solution water, supplemented with 0.05 % w/v para-amino benzoic acid (PABA), and kept at 21 °C under 70 % humidity. Mosquitoes were collected and killed in 70 % ethanol, and rinced in Leibovitz’s L-15 Medium (supplemented with fungisone, penicillin and streptomicin). Salivary glands were individually isolated 21 days after the mosquito blood meal by microdissection, and collected in DMEM medium supplemented with antibiotics. Sporozoites were released by manual grinding of salivary glands, passed on a 40 μm filter to eliminate mosquitoes large debris, and washed in DMEM. Collected sporozoites were kept at 4°C.

### Density gradient purification of P. berghei sporozoites

Density gradient purification was performed as described [5]. Briefly, a 17% w/v solution of Accudenz (Accurate Chemical #AN7050) dissolved in distilled deionized water (ddH2O) was filter sterilized and stored at 4°C. A 3 ml Accudenz cushion was loaded in a 15 ml conical tube and the dissected sporozoite mixture (up to 1 ml) was gently layered on top of the cushion. The column was spun at 2,500 *g* at 15°C for 20 minutes (no brake) and the interface was transferred to a new, clean microcentrifuge tube and spun at top speed in a microcentrifuge for three minutes. The supernatant was discarded and the pelleted sporozoites were resuspended in PBS and counted in a Neubauer chamber.

### Immunocapture of P. berghei sporozoites

Freshly dissected PbGFP sporozoites were incubated for 1 h at 4 °C in DMEM in the presence of 10 μg/ml of the anti-CSP mAb 3D11 [30]. After centrifugation, pelleted sporozoites were incubated with Protein G coupled MACS microbeads for 15 min at 4°C according to the manufacturer’s protocol. Beads were then trapped on MACS MS column and washed 5 times with PBS. Beads and sporozoites were then eluted, centrifuged and resuspended in PBS.

### Flow cytometry sorting of PbGFP sporozoites

After washing, the *PbGFP sporozoites* sorting was performed using a S3 cell sorter (BioRad). Of note, the flow sorting is performed on unfixed sporozoites in order to allow optimal proteins extraction. Fluorescent sporozoites were collected in 1X-PBS at 4°C, centrifuged (8,000g, 10 min, 4°C), and stored at −80°C until proteins extraction. Purity for PbGFP sporozoite fractions was systematically verified after sorting.

### Parasite lysis and sample preparation for mass spectrometry

Lysis of sporozoites was done directly in Laemmli buffer (62.5 mM Tris/HCl pH 6.8, 2% w/v SDS, 10% v/v glycero, 0.01% w/v Bromophenol blue and 25mM DTT). Lysates were incubated at 90 °C for 10min, centrifuged (15000g, 15min, 4°C), and stored at −80°C before analysis. Lysates were loaded on a polyacrylamide gel (SDS-PAGE) composed of a concentration gel (4% acrylamide) and a separation gel (10% acrylamide). A short electrophoresis migration was made (about 5mm). The gel was then fixed and stained with Coomassie blue Imperial Protein Stain (Thermo Fischer) for 2 hours. The short lanes containing the samples were then cut into small 1 mm^3^ pieces and placed on a 96-well pierced plate. Digestion was done by the DigestProMSi robot (Intavis). The samples were destained with a solution of 50 mM ammonium bicarbonate (AmBic) and 50% ethanol (EtOH) at 60°C. This step was followed by reduction (incubation in 10mM DTT in 50mM AmBic for 30 min at 56°C), and alkylation (incubation in 50mM iodoacetamide in 50mM AmBic for 30 min at RT in the dark). Proteins were digested with 200 ng trypsin per well overnight at 37°C. The gel pieces were washed twice in 60% acetonitrile (ACN) / 0.1% trifluoroacetic acid (TFA) for 20 minutes. Peptide extracts were then dried with Speed-Vac and resuspended in 20 μl of ACN 2% / formic acid (FA) 0.1%. Sample desalting was performed with home-made StageTips consisting in a stack of two reverse-phase C18 layers (Empore SPE Disks C18, Sigma Aldrich) inserted in a 10 μl tip. This step was carried out using the DigestProMSi robot (Intavis) and was necessary before LC-MS/MS analysis to avoid fouling of the analytical column due to residual debris. The StageTip was first hydrated with methanol and then activated by passing a 50% ACN / 0.5% acetic acid (HAc) solution and then a 0.5% HAc solution. The peptide solution previously diluted with 0.5% acetic acid to a final volume of 70 μl was then gently passed through the StageTip. The peptides retained in the StageTip were washed and desalted by passing the 0.5% HAc solution. Finally, the peptides were eluted with an ACN 80% / HAc 0.5% solution, completely dried (Speed-Vac) and resolubilized in 20 μl of ACN 2% / FA 0.1%. Samples were stored at −20 ° C until MS analysis. If necessary, peptide samples were concentrated using Speed-Vac to reduce the volume before injection into the LC-MS/MS. In some cases, we also used the pre-column (reverse phase) integrated into the LC-MS/MS system for on-line peptide desalting.

### Liquid chromatography tandem mass spectrometry

Peptides were analyzed by the new and sensitive timsTOF PRO mass spectrometer (Bruker) coupled to the nanoElute HPLC. Peptides were separated on an Odyssey column from IonOpticks (1.6μm C18, 120 Å, 75μm ID, 25 cm) using a 90 min gradient from 2 to 37 % ACN with 0.1% formic acid. MS acquisition was run in DDA mode with PASEF, from 100 to 1700 m/z with an active exclusion of 0,4 min. Parent ion selection was achieved with a two-dimensional m/z and 1/k0 selection area filter allowing exclusion of singly charged ions. Low-abundance precursors were selected several times for PASEF-MS/MS until the target value. Capillary voltage was set at 1,4 kV. The total cycle time was 1,15 sec with 10 PASEF cycles.

### Data collection and analysis

To compare our data with data from a previous proteomic analysis of *P. yoelii* sporozoites [8,10], all data were processed in the same way. Mascot generic files (mgf) were generated using Data Analysis 5.1 (Bruker) and processed with X!Tandem pipeline version 0.2.36 (http://pappso.inra.fr/bioinfo/xtandempipeline) using databases from UniProt or PlasmoDB. The databases used were: Contaminants_20160129 containing a list of common contaminants, Anopheles_stephensi_UP.fasta (Date 09/2018, number of entries: 11,767) and PlasmoDB_PB_39_PbergheiANKA (Date 09/2018, number entered: 5,076). The search parameters were: maximum 1 missed cleavage allowed, Cys-CAM as fixed modification, Met-Ox and acetyl N-term as variable modifications. MS mass error tolerances were adapted to instrument. MS1 mass error tolerance were set to 50 ppm for timsTOF PRO and 25 ppm for Orbitrap. MS2 mass error tolerance were set to 50 ppm for timsTOF PRO and 0,4 Da for Orbitrap. Protein lists were filtered and validated using Proline software version 1.6.1 (FDR < 1% at peptide and protein level, minimum 1 peptide per protein).

## Acknowledgements

We thank Jean-François Franetich, Maurel Tefit and Thierry Houpert for rearing of mosquitoes and technical assistance. This work was supported in part by the Laboratoire d’Excellence ParaFrap (ANR-11-LABX-0024), the Agence Nationale de la Recherche (ANR-16-CE15-0004 and ANR-16-CE15-0010) and the Fondation pour la Recherche Médicale (EQU201903007823). We acknowledge the Conseil Régional d’Ile-de-France, Sorbonne Université, the National Institute for Health and Medical Research (INSERM) and the Biology, Health and Agronomy Infrastructure (IBiSA) for funding the timsTOF PRO.

## Associated data

The mass spectrometry proteomics data have been deposited to the ProteomeXchange Consortium (http://proteomecentral.proteomexchange.org) via the PRIDE partner repository [31] with the dataset identifier PXD022735.

Data S1, Supporting Information lists of all proteins (1a) and peptides (1b) identified in extracts from infected mosquito salivary glands after parasite purification.

Data S2, Supporting Information lists the *P. berghei* proteins identified in purified sporozoite samples.

Data S3, Supporting Information compares the proteomes of *P. falciparum, P. vivax, P. yoelii* and *P. berghei* sporozoites.

